# LRRK2 causes centrosomal deficits via phosphorylated Rab10 and RILPL1 at centriolar subdistal appendages

**DOI:** 10.1101/2021.08.23.457380

**Authors:** Antonio Jesús Lara Ordóñez, Belén Fernández, Rachel Fasiczka, Yahaira Naaldijk, Elena Fdez, Marian Blanca Ramírez, Sébastien Phan, Daniela Boassa, Sabine Hilfiker

## Abstract

The Parkinson’s disease-associated LRRK2 kinase phosphorylates multiple Rab GTPases including Rab8 and Rab10, which enhances their binding to RILPL1 and RILPL2. The nascent interaction between phospho-Rab10 and RILPL1 blocks ciliogenesis *in vitro* and in the intact brain, and interferes with the cohesion of duplicated centrosomes in dividing cells. We show here that various LRRK2 risk variants and all currently described regulators of the LRRK2 signaling pathway converge upon causing centrosomal cohesion deficits. The cohesion deficits do not require the presence of RILPL2 or of other LRRK2 kinase substrates including Rab12, Rab35 and Rab43. Rather, they depend on the RILPL1-mediated centrosomal accumulation of phosphorylated Rab10. RILPL1 localizes to the subdistal appendages of the mother centriole, followed by recruitment of the LRRK2-phosphorylated Rab protein to cause the centrosomal defects. These data reveal a common molecular pathway by which alterations in the LRRK2 kinase activity impact upon centrosome-related events.

## Introduction

Autosomal-dominant mutations in the leucine rich repeat kinase 2 (LRRK2) gene cause familial Parkinson’s disease (PD), and coding variants in the same gene can act as risk factors for sporadic PD. Known pathogenic LRRK2 mutations produce a protein with increased kinase activity (1, 2), raising the possibility that kinase inhibitors may be useful to treat LRRK2-related PD. Therefore, understanding the downstream effects of enhanced LRRK2 kinase activity is of interest to understand disease pathomechanism(s) as well as potential therapeutics.

Mass spectroscopy efforts have revealed a subset of Rab GTPases, including Rab8, Rab10, Rab12, Rab35 and Rab43 as the primary endogenous substrates of LRRK2 (3–5). LRRK2 phosphorylates these substrates in a conserved region of the switch 2 domain, which leads to impaired interactions with various effector and regulatory proteins (3). We have previously shown that this interferes with the physiological functions of these Rab proteins as key regulators of distinct membrane trafficking events (6, 7). Importantly though, the phosphorylated Rab proteins also show enhanced binding to a set of novel effector proteins (4), raising the possibility that these nascent interactions may contribute to the pathobiology of LRRK2.

LRRK2-phosphorylated Rab8 and Rab10 bind with great preference to RILPL1 and RILPL2 (4), two poorly characterized proteins reported to regulate ciliary content (8). An important and direct consequence of the LRRK2-mediated phosphorylation of these Rab proteins is a decrease in primary cilia in various cell types *in vitro* as well as in the intact mouse brain (4, 9, 10). We previously showed that pathogenic LRRK2 not only causes ciliary deficits, but also deficits in the cohesion between duplicated centrosomes in a manner mediated by the phosphorylated Rab proteins and by RILPL1 (10, 11). Such centrosomal cohesion deficits can also be observed in peripheral cells derived from LRRK2 PD patients as compared to healthy controls (12). Thus, ciliary deficits in brain, and centrosomal cohesion deficits in dividing cells are distinct cellular reflections of the same LRRK2-mediated phospho-Rab/RILPL1 interaction.

Recent studies have described upstream regulators of the LRRK2 kinase activity including Rab29 and vps35. Rab29 is a protein which modulates PD risk (13), and under certain conditions is able to stimulate the LRRK2 kinase activity (14–16). A point mutation in vps35, the cargo binding component of the retromer complex, causes autosomal-dominant late-onset familial PD (17–19), and potently activates the LRRK2 kinase activity as assessed by Rab10 phosphorylation in cells and tissues (20). Conversely, the PPM1H phosphatase acts as a downstream regulator to counteract LRRK2 signaling by selectively dephosphorylating Rab8 and Rab10 (21). Finally, LRRK2 harbours several protein coding variants which modulate risk for sporadic PD, potentially mediated by subtle alterations in the LRRK2 kinase activity (2).

Here, we show that distinct modulators of the LRRK2 signaling pathway converge upon causing centrosomal cohesion deficits in cultured cells. The pathogenic LRRK2-mediated centrosomal cohesion deficits are independent of the presence of Rab12, Rab35, Rab43 or RILPL2, but depend on the RILPL1-mediated centrosomal accumulation of phosphorylated Rab10. Correlated light and electron microscopy (CLEM) indicates that RILPL1 directs the phosphorylated Rab proteins to subdistal appendages of the mother centriole, where they may interfere with proper centrosomal cohesion and ciliogenesis as observed in the context of pathogenic LRRK2.

## Results

### LRRK2 risk variants modulate centrosomal cohesion in HEK293T cells

To explore the relationship between centrosomal cohesion phenotypes and LRRK2 variants described to positively or negatively impact PD risk, we transiently transfected HEK293T cells with wildtype LRRK2, with a point mutant described to be non-pathogenic (T1410A), distinct point mutants described to increase PD risk (R1628P, S1647T, N2081D, G2385R), or a pathogenic point mutant which served as a positive control (Y1699C) (22–28). Compared to expression of wildtype LRRK2, the pathogenic Y1699C-LRRK2 mutant caused a pronounced deficit in centrosomal cohesion, as evidenced by the percentage of cells displaying duplicated centrosomes with a distance further than 1.5 microns apart, which was significantly reduced by transient application of the LRRK2 kinase inhibitor MLi2 (**Figure 1A,B**). Expression of the non-pathogenic T1410A LRRK2 mutant was without effect, whilst four distinct PD risk variants caused a statistically significant centrosomal cohesion deficit which was reverted by MLi2 (**Figure 1A,B**). As assessed by immunoblotting, transient expression of the pathogenic Y1699C-LRRK2 mutant caused a detectable increase in Rab10 phosphorylation as compared to wildtype LRRK2, whilst no detectable differences were observed when expressing the various LRRK2 risk variants (**Figure 1C**). However, increased accumulation of phospho-Rab10 in individually transfected cells could be detected by immunocytochemistry when expressing pathogenic LRRK2 or the various risk variants, and such accumulation was reverted in all cases by LRRK2 kinase inhibitor (**Figure 1D**).

**Figure 1.**
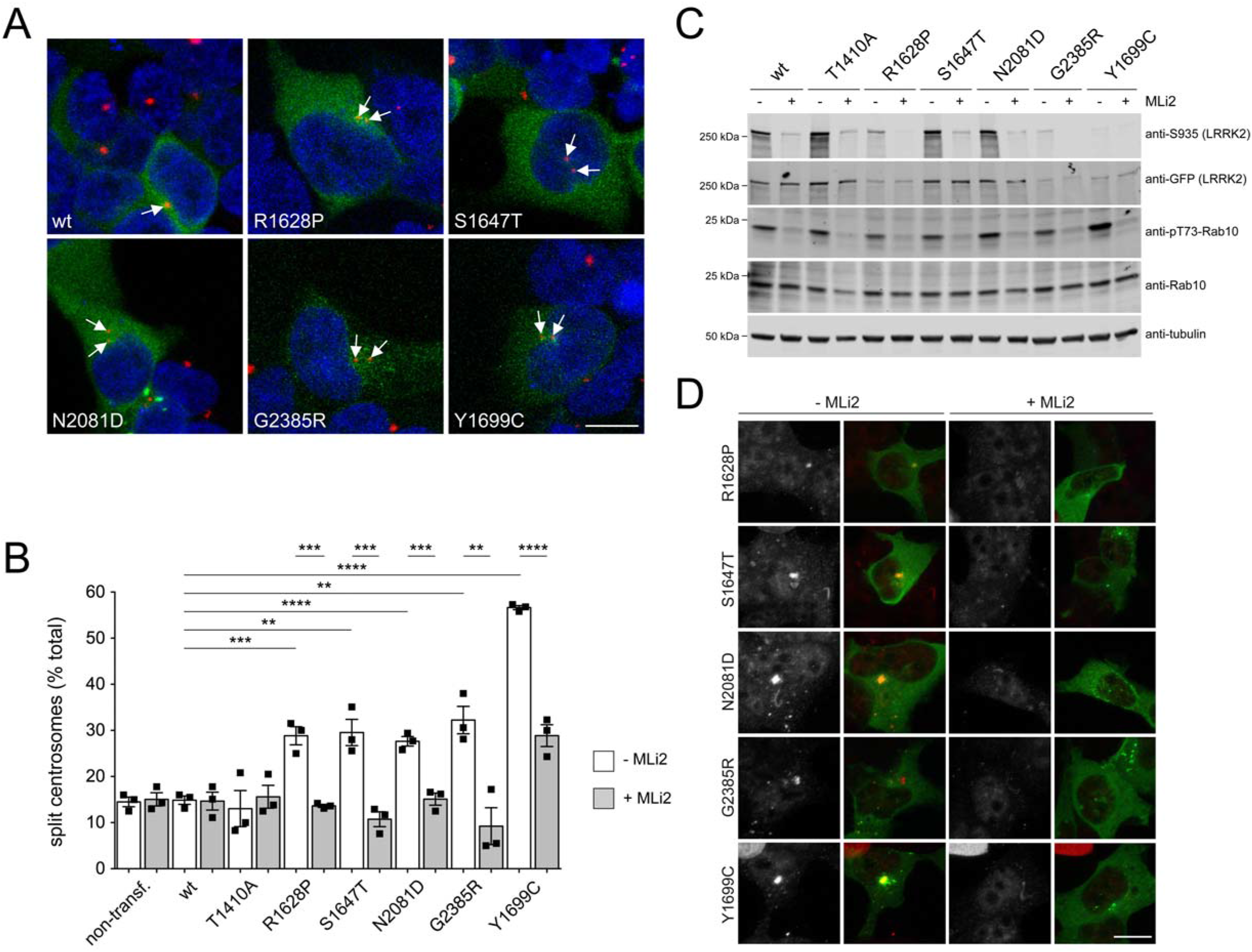
LRRK2 risk variants cause centrosomal cohesion deficits. (**A**) HEK293T cells were transfected with GFP-tagged wildtype (wt) LRRK2, risk variants as indicated, or with pathogenic Y1699C LRRK2. Cells were stained for the centrosomal marker pericentrin (red) and DAPI (blue). Arrows point to transfected cells with duplicated centrosomes further than 1.5 microns apart (split centrosomes). Scale bar, 10 μm. (**B**) Quantification of the percentage of cells with duplicated split centrosomes from either non-transfected (ctrl) cells, or from cells transfected with the indicated constructs, in the absence or presence of MLi2 (100 nM, 2 h) as indicated. Bars represent mean ± S.E.M. (n=3 experiments); ****p < 0.001; ***p < 0.005; **p < 0.01. (**C**) Cells were transfected with the indicated constructs, left untreated or treated with MLi2 (100 nM, 2 h) as indicated, and extracts blotted for GFP-tagged LRRK2, phosphorylated LRRK2 (S935), phosphorylated Rab10 (pT73-Rab10), total Rab10, and tubulin as loading control. (**D**) Cells were transfected with the indicated GFP-tagged constructs, left untreated or treated with MLi2 as indicated, and stained with an antibody against endogenous phosphorylated Rab10 (red). Scale bar, 10 μm.

To analyse the effect of the R1398H mutation in LRRK2 which is protective against PD (25, 29, 30), we introduced it into either wildtype or pathogenic LRRK2 constructs. Transient expression of pathogenic G2019S, R1441C, Y1699C, N1437H or I2020T LRRK2 mutants caused a significant deficit in centrosomal cohesion which was attenuated by introduction of the protective R1398H variant in all cases (**Figure S1**). Similar results were obtained when introducing synthetic mutations (R1398L or R1398L/T1343V) described to alter Rab10 phosphorylation by modulating LRRK2 GTP binding/hydrolysis (31–34) (**Figure S2**). Collectively, these data indicate that risk or protective LRRK2 variants can either negatively or positively impact upon the centrosomal cohesion deficits mediated by the LRRK2 kinase activity.

### Rab12, Rab35 and Rab43 are not required for the centrosomal cohesion deficits mediated by pathogenic LRRK2

LRRK2 phosphorylates various endogenous Rab proteins including Rab8, Rab10, Rab12, Rab35 and Rab43 (4), and previous studies have implicated Rab8 and Rab10 in the ciliary and centrosomal deficits mediated by pathogenic LRRK2 (9, 10). To study the potential role of Rab12, Rab35 or Rab43, we employed A549 cells where these proteins were knocked out using CRISPR-Cas9 (4) (**Figure 2A,B**). Consistent with what we have previously shown (10), expression of pathogenic LRRK2 in wildtype A549 cells caused a centrosomal cohesion deficit which was reverted by MLi2, whilst wildtype or a kinase-inactive LRRK2 mutant were without effect (**Figure 2C**). Similar centrosomal cohesion deficits were observed when pathogenic LRRK2 constructs were expressed in A549 cells deficient in either Rab12 (**Figure 2B,D**), Rab35 (**Figure 2E**) or Rab43 (**Figure 2F**). In all cases, the deficits were reverted by MLi2, and were not observed when expressing wildtype or kinase-inactive LRRK2 mutant. The pathogenic LRRK2-mediated increase in the percentage of split centrosomes was paralleled by an increase in the overall mean distance between duplicated centrosomes (**Figure 2G**). Furthermore, and as assessed by immunoblot analysis, expression of pathogenic LRRK2 caused similar increases in the levels of phospho-Rab8 and phospho-Rab10 in wildtype cells or in cells deficient in either Rab12, Rab35 or Rab43 (**Figure 2H**). Thus, and in contrast to Rab8 and Rab10 (10), the centrosomal cohesion deficits mediated by pathogenic LRRK2 are not dependent on the presence of Rab12, Rab35 or Rab43.

**Figure 2.**
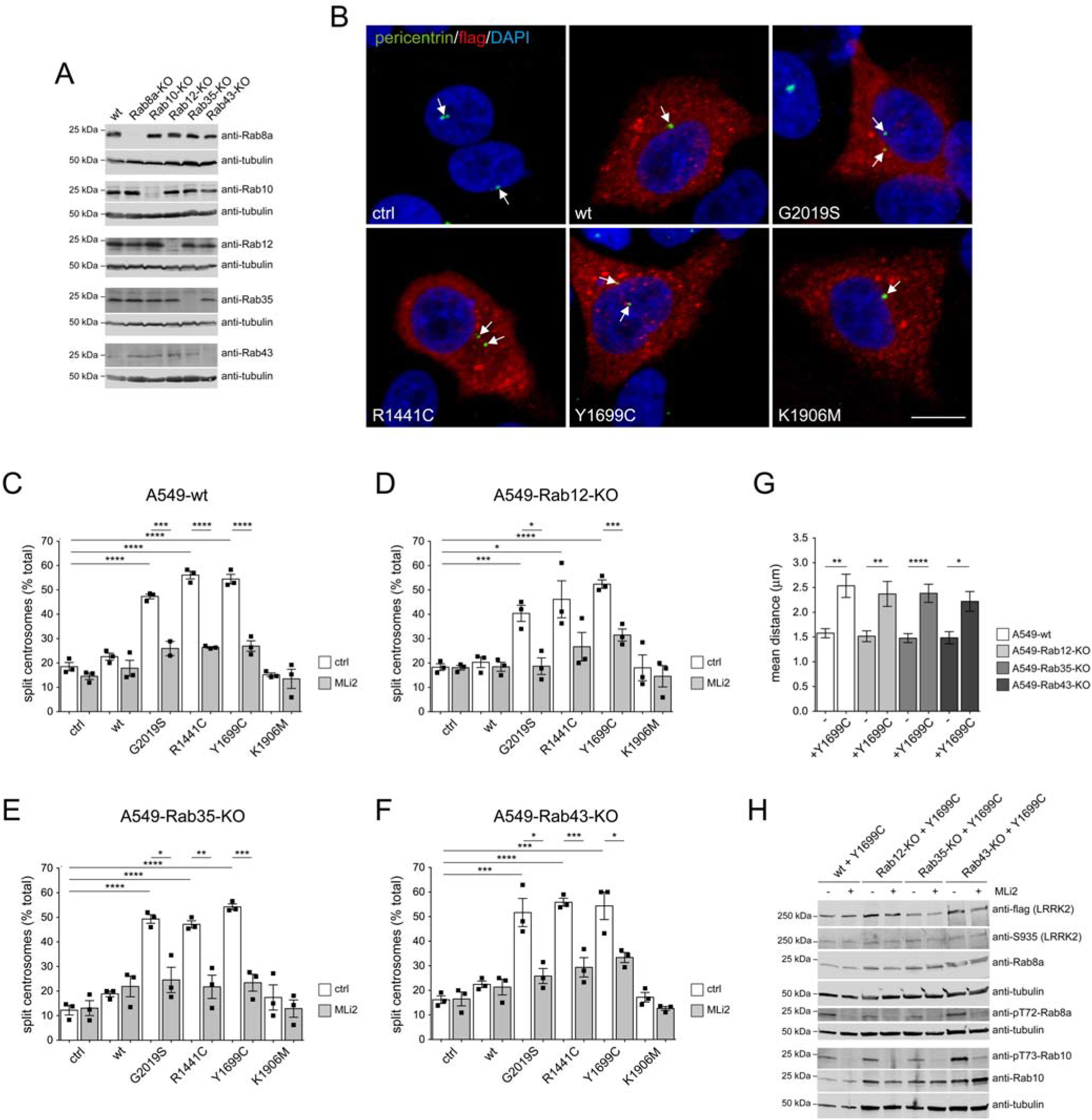
Pathogenic LRRK2-mediated centrosomal cohesion deficits do not depend on the presence of Rab12, Rab35 or Rab43. (**A**) Wildtype A549 cells (wt), or cells where the distinct Rab proteins had been knocked out (KO) using CRISPR-Cas9 were subjected to immunoblot analysis for the presence or absence of the various Rab proteins as indicated, with tubulin as loading control. (**B**) Example of A549 Rab12-KO cells transfected with either pCMV (ctrl), or with the indicated flag-tagged LRRK2 constructs and stained with antibodies against flag, pericentrin and with DAPI. Scale bar, 10 μm. (**C**) Quantification of the percentage of wildtype A549 cells with split centrosomes (duplicated centrosomes with a distance between their centres > 2.5 μm) transfected with the different LRRK2 constructs, and either left untreated or incubated with 500 nM MLi2 for 2 h prior to immunocytochemistry as indicated. (**D**) Same as in (C), but employing Rab12-KO cells. (**E**) Same as in (C), but employing Rab35-KO cells. (**F**) Same as in (C), but employing Rab43-KO cells. In all cases, bars represent mean ± S.E.M. (n=3 experiments); ****p < 0.001; ***p < 0.005; **p < 0.01; *p < 0.05. (**G**) Wildtype cells, or Rab12-KO, Rab35-KO or Rab43-KO cells were transfected with flag-tagged Y1699C LRRK2, and distances between duplicated centrosomes quantified from around 50-70 transfected or non-transfected cells each. ****p < 0.001; **p < 0.01; *p < 0.05. (**H**) A549 wt cells, or Rab12-KO, Rab35-KO or Rab43-KO cells were transfected with Y1699C-mutant LRRK2 construct, left untreated or incubated with 500 nM MLi2 for 2 h as indicated, and extracts were blotted for flag-tagged LRRK2, phosphorylated LRRK2 (S935), pT72-Rab8a, total Rab8a, pT73-Rab10, total Rab10, or tubulin as loading control.

### Upstream and downstream regulators of the LRRK2 signaling pathway impact upon centrosomal cohesion

Previous work has indicated that co-expression of Rab29 and LRRK2 stimulates the LRRK2 activity by recruiting it to the Golgi complex (14, 15), and our data indicate that this is associated with centrosomal cohesion deficits (35). To determine whether endogenous Rab29 is required for the pathogenic LRRK2-mediated centrosomal deficits, we employed A549 cells deficient in Rab29. The centrosomal cohesion deficits mediated by pathogenic LRRK2 expression were significantly blunted in Rab29-deficient as compared to wildtype cells (**Figure S3**). Since knockout of Rab29 does not seem to influence basal or pathogenic LRRK2 kinase activity (16), these findings suggest that the presence of Rab29 may be important for the LRRK2-mediated centrosomal cohesion phenotype in a manner independent of its ability to regulate the LRRK2 kinase activity.

Vps35 is a key component of the retromer complex which regulates vesicular trafficking to and from the Golgi complex. Strikingly, a point mutation (vps35-D620N) which causes autosomal-dominant late-onset PD (17–19) hyperactivates LRRK2 through a currently unknown mechanism (20). We next wondered whether vps35 or mutants thereof may impact upon the LRRK2-mediated centrosomal cohesion deficits. Pathogenic LRRK2 expression in vps35-deficient A549 cells (20) caused centrosomal cohesion deficits identical to those observed in wildtype cells, indicating that the presence of endogenous vsp35 is not required for this phenotype (**Figure S3**). To determine whether the pathogenic vps35-D620N mutant regulates the centrosomal cohesion deficits by activating LRRK2, we coexpressed wildtype LRRK2 with HA-tagged wildtype or mutant vps35 variants in HEK293T cells. Coexpression of LRRK2 with vsp35-D620N caused a centrosomal cohesion deficit which was reverted by MLi2 treatment (**Figure 3A,B**) and correlated with an increase in the levels of phospho-Rab10 as assessed by immunoblot analysis (**Figure 3C**). In contrast, no effects were observed when coexpressing wildtype LRRK2 with either wildtype vps35, or with two distinct, non-pathogenic vps35 point mutants (vps35-L774M, vps35-M57I) (17, 19) (**Figure 3A-C**). These data support the conclusion that the vps35-D620N mutant activates the LRRK2 kinase as assessed by increased Rab10 phosphorylation to cause centrosomal cohesion deficits.

**Figure 3.**
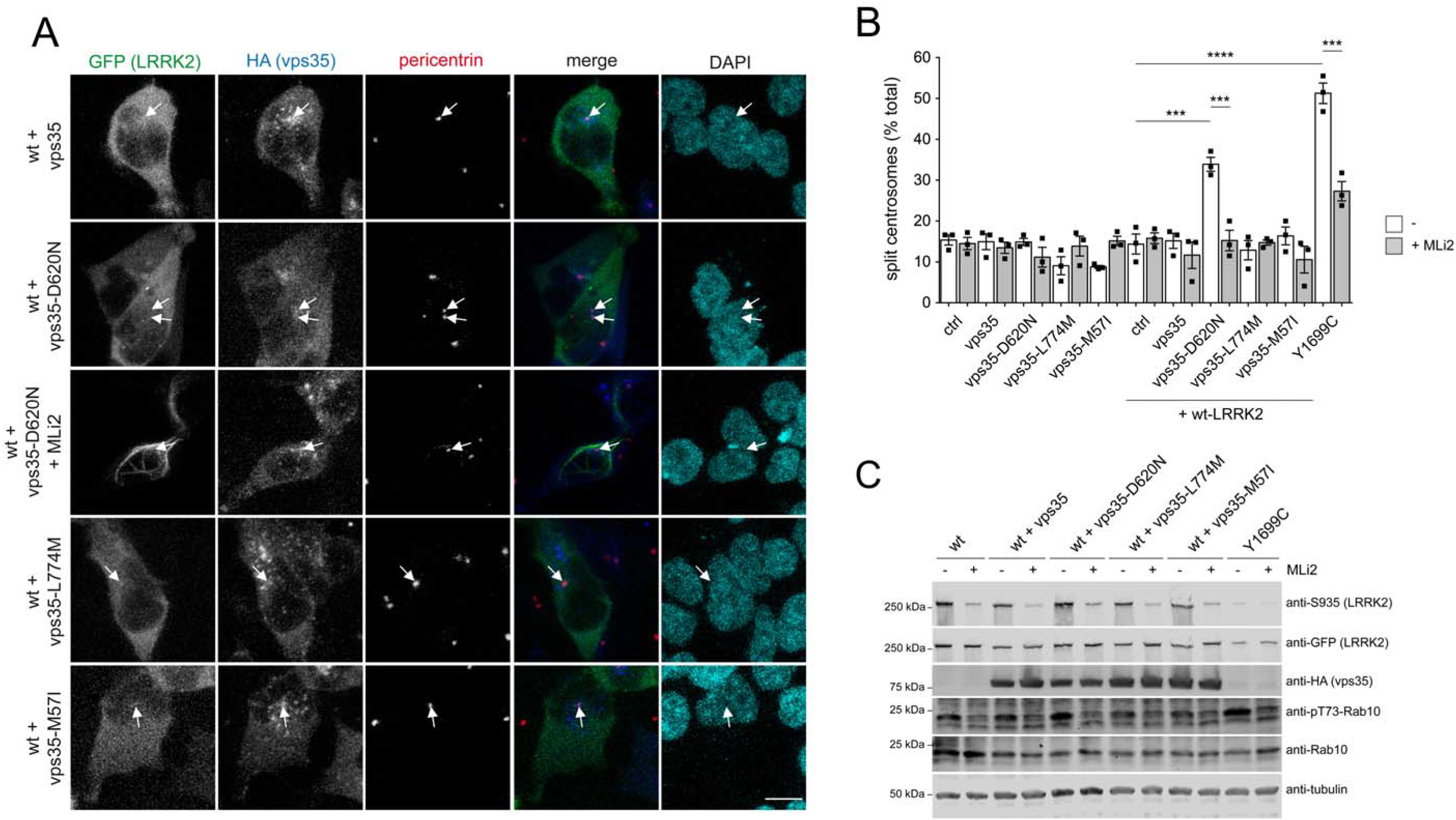
Vps35-D620N expression with LRRK2 causes centrosomal cohesion deficits dependent on the LRRK2 kinase activity. (A) HEK293T cells transfected with GFP-tagged wildtype LRRK2 (wt) and HA-tagged wildtype or mutant vps35 constructs, and treated with or without MLi2 (100 nM, 2 h) prior to immunocytochemistry as indicated. Cells were stained with an antibody against the HA-tag (Alexa-647 secondary antibody; pseudocolored in blue), an antibody against pericentrin (Alexa-555 secondary antibody; red) and DAPI (cyan). Scale bar, 10 μm. (**B**) Quantification of the percentage of cells with duplicated split centrosomes transfected with pCMV (ctrl) or different HA-tagged vps35 constructs, or co-transfected with GFP-tagged wildtype LRRK2, in the absence or presence of MLi2 (100 nM, 2 h) as indicated. Bars represent mean ± S.E.M. (n=3 experiments); ****p < 0.001; ***p < 0.005.

Lastly, we evaluated the contribution of PPM1H, the protein phosphatase responsible for dephosphorylating the LRRK2-phosphorylated Rab10 and Rab8 proteins (21). Side-by-side comparison revealed that wildtype LRRK2 expression *per se* was able to cause a centrosomal cohesion deficit in the PPM1H knockout cells, but not in wildtype cells (**Figure 4A,B**). The centrosomal deficits were reverted by MLi2, and correlated with an increase in the levels of phospho-Rab10 and phospho-Rab8 as assessed by immunoblotting (**Figure 4C**). No additional effects on centrosomal cohesion were observed when expressing distinct pathogenic LRRK2 mutants in the PPM1H knockout as compared to the control cells (**Figure 4C**). Therefore, altogether these data support the conclusion that the centrosomal cohesion deficits mediated by the LRRK2 kinase activity are subject to modulation by both upstream and downstream components of the LRRK2 signaling pathway.

**Figure 4.**
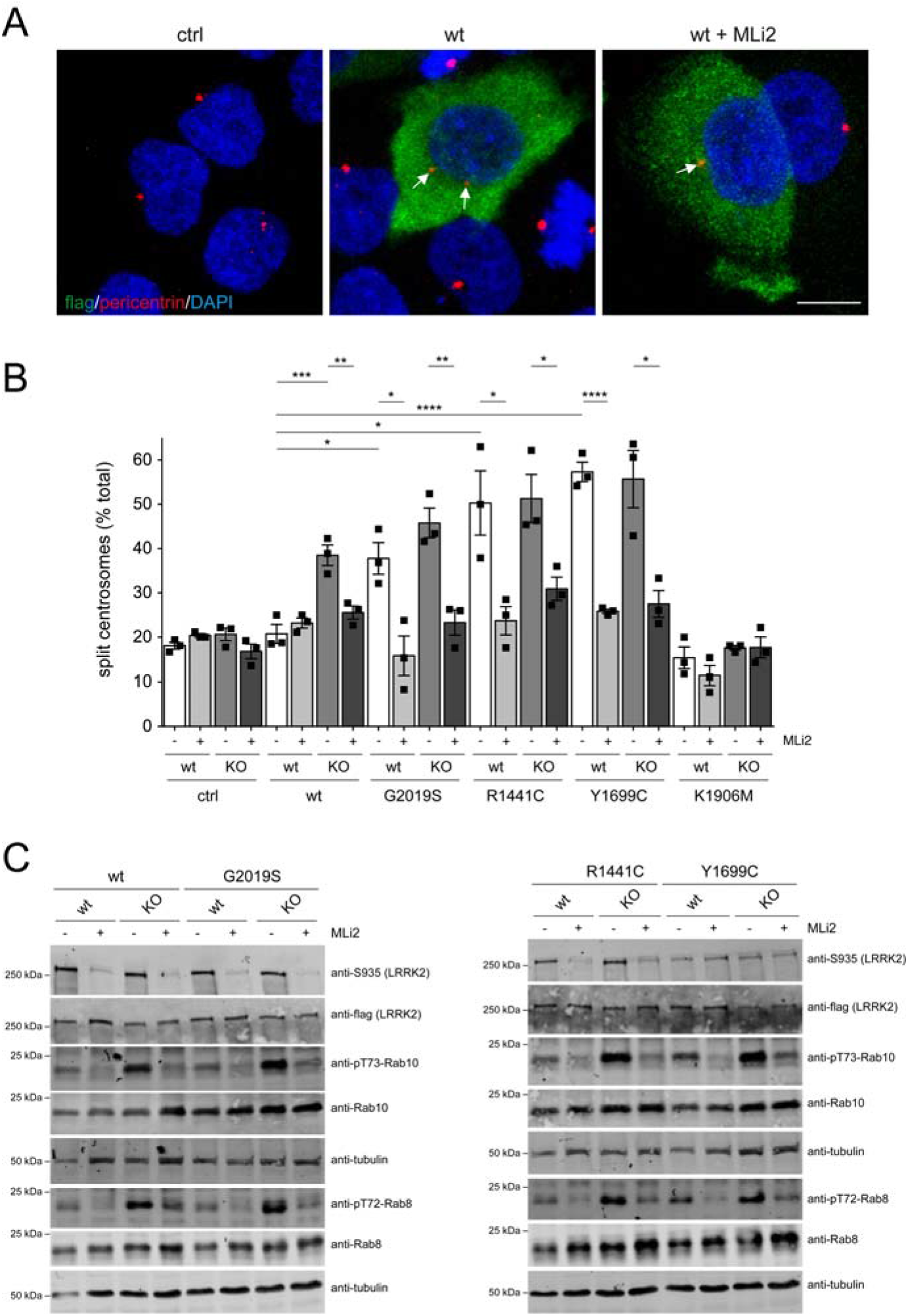
Wildtype LRRK2 expression causes centrosomal cohesion deficits in PPM1H knock-out A549 cells. (**A**) Example of PPM1H-KO cells transfected with pCMV (ctrl) or with flag-tagged wildtype LRRK2, in the presence or absence of MLi2 (500 nM, 2 h) as indicated before immunocytochemistry with antibody against flag (green), pericentrin (red) and DAPI. Scale bar, 10 μm. (**B**) Quantification of the percentage of A549 wildtype or PPM1H-KO cells displaying split centrosomes (duplicated centrosomes with a distance between their centres > 2.5 μm) transfected with either pCMV (ctrl) or the different LRRK2 constructs, and either left untreated or incubated with MLi2 (500 nM, 2 h) prior to immunocytochemistry. Bars represent mean ± S.E.M. (n=3 experiments); ****p < 0.001; ***p < 0.005; **p < 0.01; *p < 0.05. (**C**) A549 wildtype or PPM1H-KO cells were transfected with the indicated LRRK2 constructs, left untreated or incubated with MLi2 (500 nM, 2h) as indicated, and extracts were blotted for flag-tagged LRRK2, phosphorylated LRRK2 (S935), pT72-Rab8a, total Rab8a, pT73-Rab10, total Rab10, or tubulin as loading control.

### RILPL1 localization is crucial for the centrosomal cohesion deficits mediated by LRRK2-phosphorylated Rab proteins

LRRK2-phosphorylated Rab8 and Rab10 bind with strong preference to RILPL1 and RILPL2 (4), and RILPL1 is required for the centrosomal cohesion deficits mediated by pathogenic LRRK2 (10). To evaluate the involvement of RILPL2, we transfected A549 RILPL2 knockout cells with wildtype or distinct pathogenic LRRK2 mutants. Pathogenic LRRK2 expression caused centrosomal cohesion deficits in RILPL2 knockout cells identical to those observed in wildtype cells, which were reverted by MLi2 in all cases (**Figure 5**). In addition, wildtype cells transfected with pathogenic LRRK2 displayed prominent phospho-Rab10 staining in a perinuclear area and in tubular structures, and similar staining was observed in RILPL2 knockout cells (**Figure S4**). In contrast, and as previously reported (9), RILPL1 knockout cells transfected with pathogenic LRRK2 showed a diminished perinuclear distribution of phospho-Rab10 accompanied by punctate staining throughout the cytosol (**Figure S4**). Together, these data support the conclusion that the centrosomal cohesion phenotype mediated by pathogenic LRRK2 is largely determined by the interaction of phospho-Rab10 with RILPL1, at least in this cell system.

**Figure 5.**
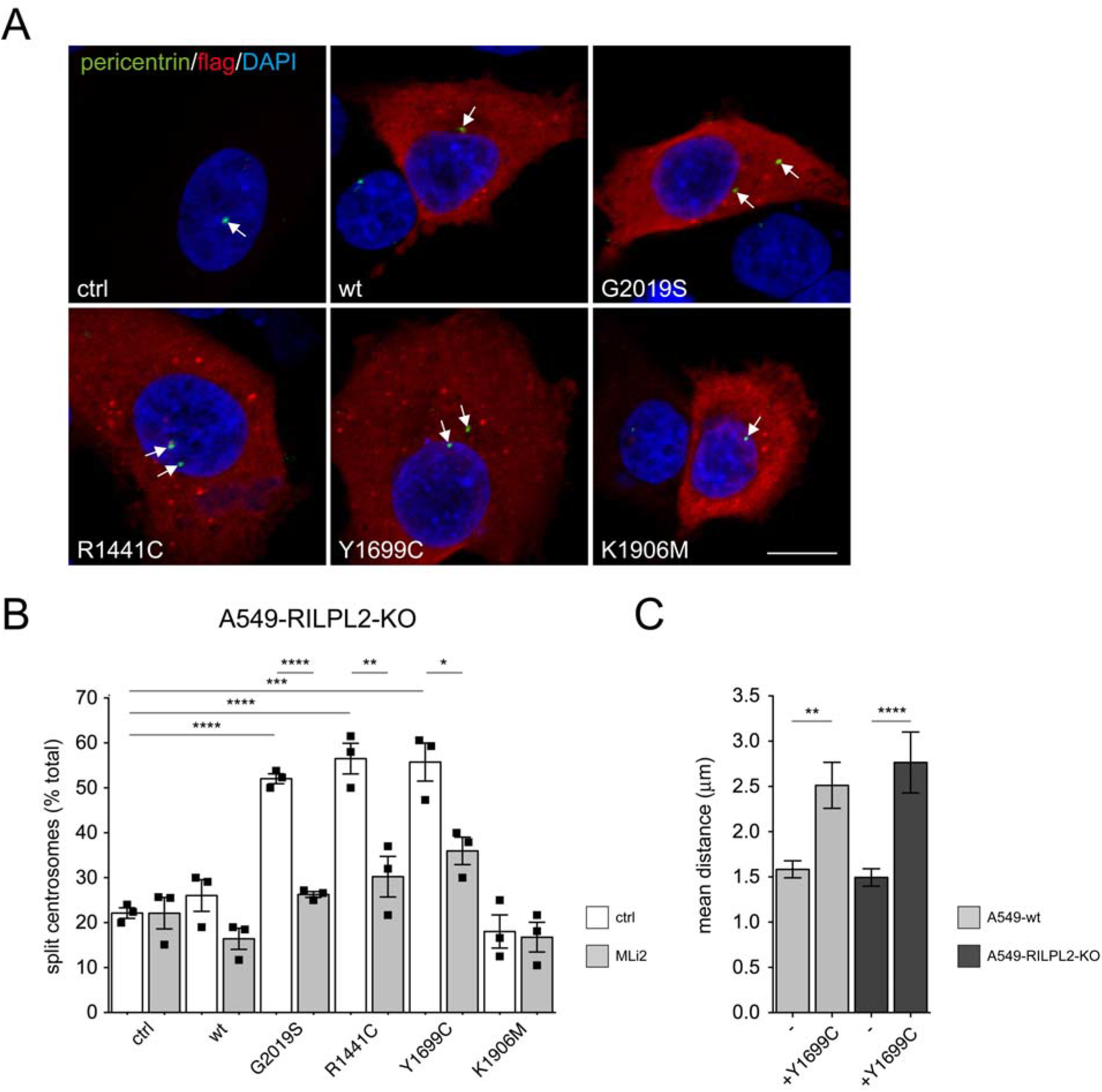
RILPL2 is dispensable for the LRRK2-mediated centrosomal cohesion deficits. (**A**) RILPL2-KO cells were transfected with either pCMV (ctrl), or with the indicated flag-tagged LRRK2 constructs, and stained with antibodies against flag, pericentrin and with DAPI. Scale bar, 10 μm. (**B**) Quantification of the percentage of A549 RILPL2-KO cells with split centrosomes upon transfection of the indicated constructs, in either the absence or presence of MLi2 (500 nM, 2 h) prior to immunocytochemistry. Bars represent mean ± S.E.M. (n=3 experiments); ****p < 0.001; ***p < 0.005; **p < 0.01; *p < 0.05. (**C**) Wildtype or RILPL2-KO cells were transfected with flag-tagged Y1699C LRRK2, and distances between duplicated centrosomes quantified from around 50-70 transfected or non-transfected cells each. ****p < 0.001; **p < 0.01.

To further study the involvement of RILPL1 in the LRRK2-mediated centrosomal cohesion deficits, we expressed the C-terminal half of the protein (RL1d-GFP), reported to be responsible for its interaction with the phosphorylated Rab proteins (4, 9). When expressed in A549 cells, RL1d-GFP displayed a punctate as well as cytosolic localization (**Figure 6A**). Strikingly, RL1d-GFP expression completely reverted the centrosomal cohesion phenotype induced by pathogenic LRRK2, whilst not displaying an effect when expressed on its own (**Figure 6A-C**). Co-expression of RL1d-GFP with pathogenic LRRK2 did not decrease the total levels of phospho-Rab8 or phospho-Rab10 as assessed by immunoblot analysis (**Figure 6D**). Rather, whilst pathogenic LRRK2 caused a pronounced perinuclear accumulation of phospho-Rab10, co-expression with RL1d-GFP caused the redistribution of phospho-Rab10 to cytosolic RL1d-GFP-positive punctae (**Figure 6E**). These data indicate that the centrosomal phospho-Rab10 accumulation depends on the centrosomal localization of RILPL1 which is required to cause the cohesion deficits mediated by pathogenic LRRK2.

**Figure 6.**
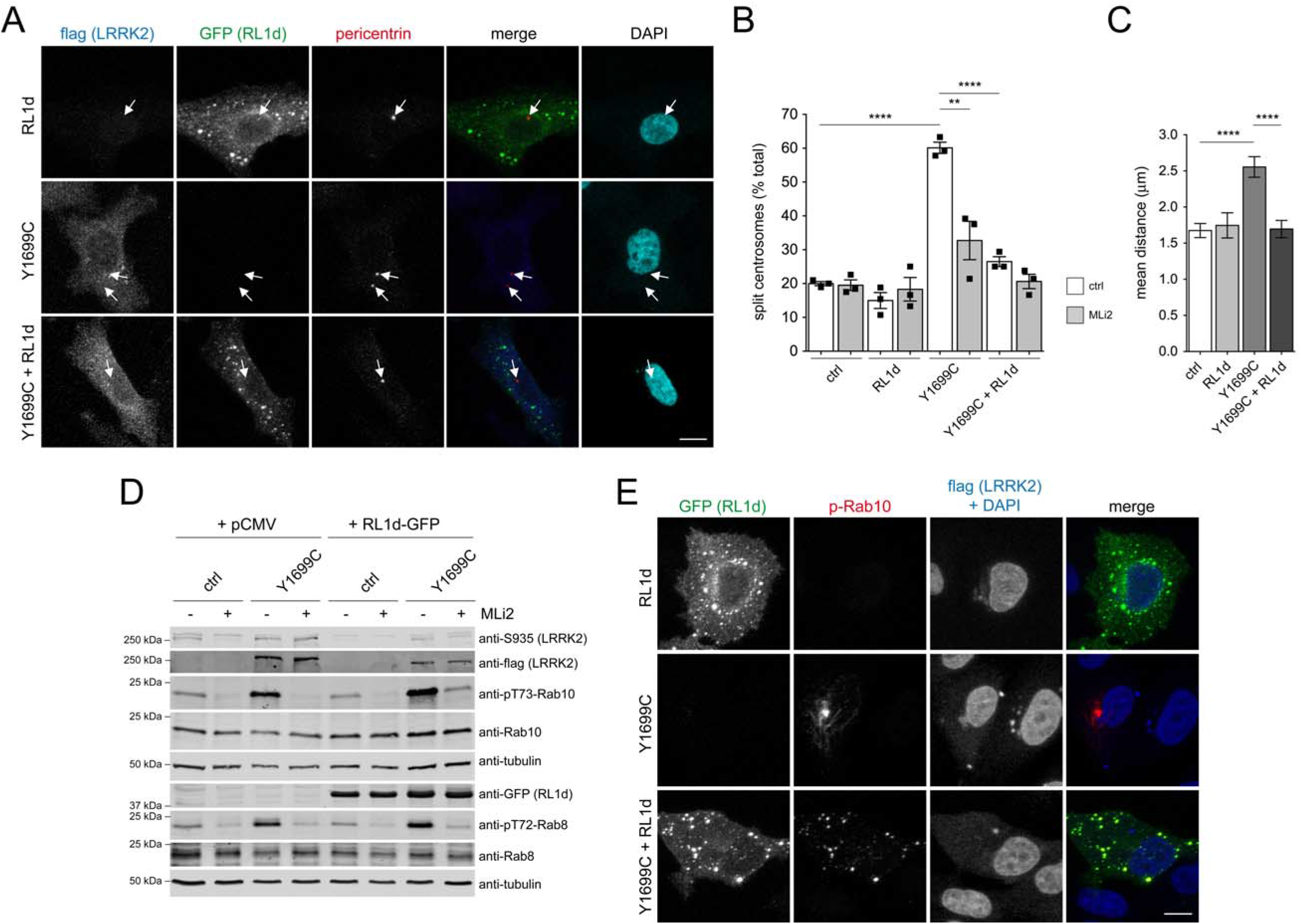
Expression of C-terminal RILPL1 reverts the LRRK2-mediated centrosomal cohesion deficits and redistributes phospho-Rab10. (**A**) Example of A549 cells transfected with the C-terminal region of RILPL1 (RL1d-GFP), flag-tagged Y1699C mutant LRRK2, or both as indicated, and stained with antibodies against flag (Alexa-594 secondary antibody; pseudocolored in blue), pericentrin (Alexa-647 secondary antibody; red) and with DAPI. Scale bar, 10 μm. (**B**) Quantification of the percentage of cells displaying split centrosomes (duplicated centrosomes with a distance between their centres > 2.5 μm) transfected with either pCMV (ctrl), RL1d-GFP, flag-tagged Y1699C LRRK2, or with both, and either left untreated or incubated with MLi2 (500 nM, 2 h) prior to immunocytochemistry. Bars represent mean ± S.E.M. (n=3 experiments); ****p < 0.001; **p < 0.01. (**C**) Cells transfected with the indicated constructs were processed for immunocytochemistry, and the distances between duplicated centrosomes were quantified from around 50 cells each. ****p < 0.001. (**D**) Cells were co-transfected with the indicated constructs, left untreated or incubated with MLi2 (500 nM, 2h) as indicated, and extracts were blotted for flag-tagged LRRK2, phosphorylated LRRK2 (S935), pT72-Rab8a, total Rab8a, pT73-Rab10, total Rab10, or tubulin as loading control. (**E**) Cells were transfected with the indicated constructs and stained with antibodies against phospho-Rab10 (Alexa-594 secondary antibody, red), flag (Alexa-405 secondary antibody, blue) and DAPI. Scale bar, 10 μm.

### RILPL1 localizes to subdistal appendages of the mother centriole

RILPL1 associates with the mother centriole which becomes the basal body that nucleates the primary cilium (8). Employing two distinct anti-RILPL1 antibodies in HEK293T cells, we corroborated the centrosomal localization of endogenous RILPL1, and the colocalization of RILPL1 with endogenous phosphorylated Rab proteins in cells transfected with pathogenic LRRK2 (**Figure S5**). Both N-terminally or C-terminally GFP-tagged RILPL1 proteins displayed a pericentrosomal localization when expressed in A549 cells (9) (**Figure 7A**), and an identical pericentrosomal localization was observed when RILPL1 was fused to miniSOG (RILPL1-miniSOG), a tag suitable for correlative light and electron microscopy (CLEM) (36, 37). In addition, and as previously reported (10), transient expression of tagged RILPL1 in A549 cells had no effect on centrosomal cohesion as assessed by measuring the mean distance between duplicated centrosomes (**Figure 7B**).

**Figure 7.**
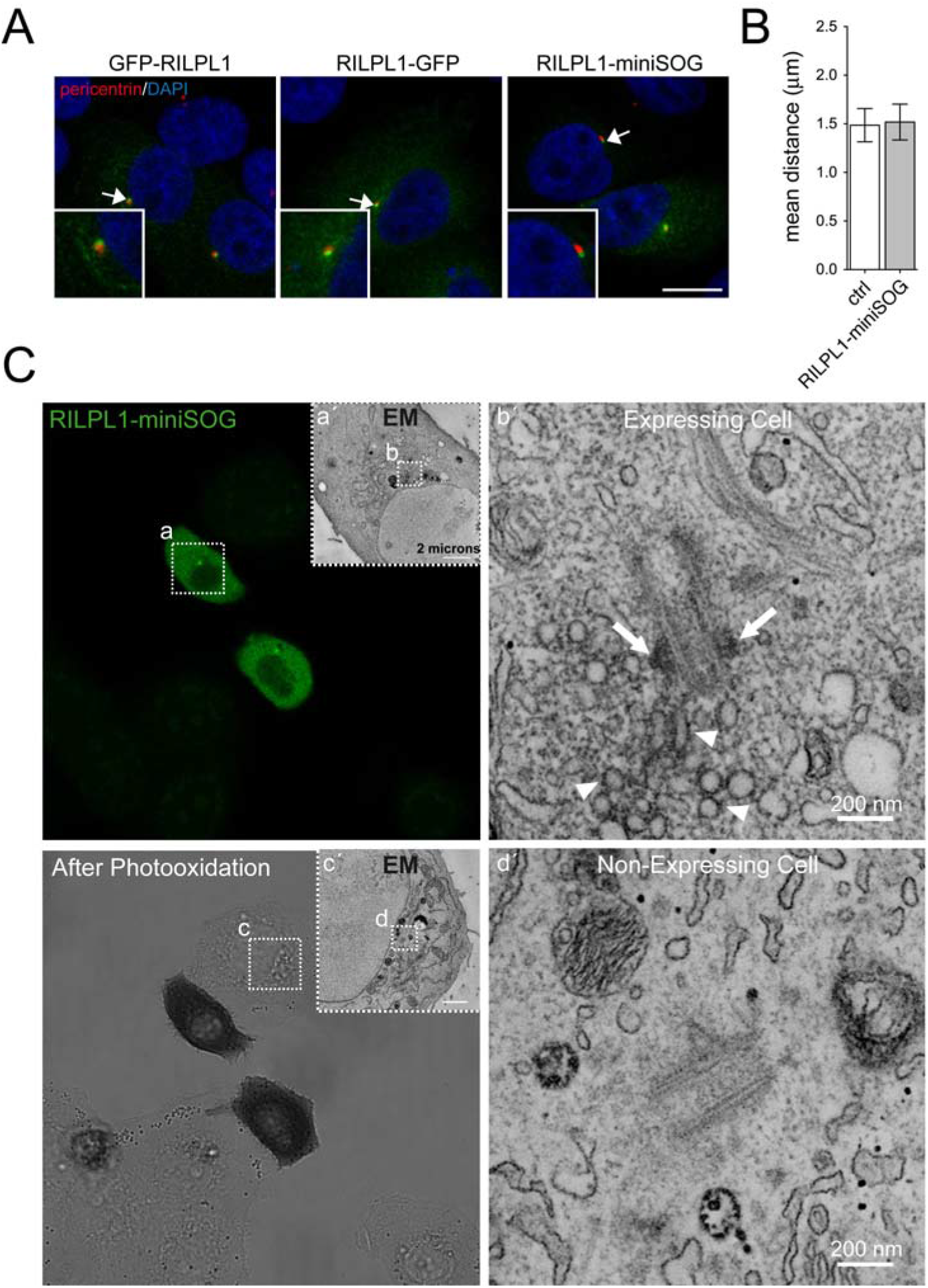
RILPL1 localizes to subdistal appendages of the mother centriole. (**A**) Example of A549 cells transfected with GFP-RILPL1, RILPL1-GFP or RILPL1-miniSOG, and stained with antibody against pericentrin (red) and with DAPI. Scale bar, 10 μm. (**B**) A549 cells were transfected with RILPL1-miniSOG and processed for immunocytochemistry, and the distances between duplicated centrosomes quantified from around 30 non-transfected and transfected cells. Bars represent mean ± S.E.M. **(C)**Correlated light and EM imaging of RILPL1-miniSOG. Transfected A549 cells were revealed first by confocal fluorescence and then by transmitted light imaging following DAB photooxidation, where an optically dense reaction product was observed in the two expressing cells. To better discriminate the DAB precipitate, the tannic acid and uranyl acetate stainings were omitted. Low magnification TEM image of the RILPL1-miniSOG-expressing cell (**a’**) corresponds to the same area indicated by the white square (**a**); similarly, the high magnification image (**b’**) corresponds to the same area indicated by the white square (**b**). White arrows point at DAB-labeled subdistal appendages of the mother centriole; arrowheads point at DAB-labeled preicentrosomal vesicles. As comparison, the low magnification TEM image of the non-expressing cell (**c’**) corresponds to the same area indicated by the white square (**c**); the high magnification image (**d’**) corresponds to the same area indicated by the white square (**d**). No labeling was observed at the subdistal appendages in the control, non-expressing cell. These observations were confirmed in cells (RILPL1-miniSOG-expressing and controls, non-expressing) from 3 different areas.

In order to image the pericentrosomal localization of RILPL1 with high spatial resolution, we performed miniSOG-induced DAB oxidation which generated a localized polymeric precipitate that could be readily identified by CLEM (**Figure 7C**). Using STEM tomography (for which we combined a multiple-tilt tomography approach with the scanning mode of a TEM), we determined that RILPL1 was localized to the subdistal appendages of the mother centriole, hinting that the interaction between phospho-Rab8/phospho-Rab10 and RILPL1 may occur at this localization. The specific DAB labelling was clearly distinguishable in RILPL1-miniSOG-expressing A549 cells as compared to adjacent non-expressing cells that were exposed to the same processing within the photooxidized area. An accumulation of DAB-labeled pericentrosomal vesicles was observed in expressing cells only, suggesting this might be induced by RILPL1 overexpression.

## Discussion

Pathogenic LRRK2 activity causes loss of primary cilia in various cell types *in vitro*, and the intact brain of G2019S or R1441C LRRK2 knockin mice *in vivo* (9, 38). Conversely, in dividing cells pathogenic LRRK2 activity causes a deficit in the proper cohesion of duplicated centrosomes (10–12). Both ciliogenesis and centrosomal cohesion deficits are due to an accumulation of LRRK2-phosphorylated Rab proteins at a centrosomal location which is dependent on RILPL1 (9, 10). In this study, we show that centrosomal cohesion is also regulated by risk and protective LRRK2 variants, and by modulators of the LRRK2 kinase signaling pathway. Expression of LRRK2 risk variants causes centrosomal cohesion deficits, whilst introduction of a protective variant into distinct pathogenic LRRK2 constructs decreases such deficits. Moreover, a point mutation in vps35 (vps35-D620N) which causes autosomal-dominant PD (17–19) and activates the LRRK2 kinase (20) causes a pronounced centrosomal cohesion deficit. Knockout of PPM1H, the phosphatase specific for LRRK2-phosphorylated Rab proteins (21) impairs centrosomal cohesion in the presence of LRRK2 expression, and a recent study shows that heterozygous loss of the PPM1H phosphatase impairs ciliogenesis in the intact mouse brain (38). These data provide strong evidence for the importance of the LRRK2 kinase activity in regulating both ciliogenesis and centrosomal cohesion deficits in non-dividing and dividing cells, respectively. In addition, we find that the centrosomal defects require accumulation of phosphorylated Rab10 in a manner dependent on RILPL1, which is localized to the subdistal appendages of the mother centriole. These data strengthen the importance of RILPL1 as a key player for the LRRK2-mediated deficits by enabling recruitment of phosphorylated Rab proteins to the mother centriole, followed by downstream events which culminate in centrosomal and ciliogenesis defects.

We find that all currently described modulators of the LRRK2 kinase activity impact upon centrosomal cohesion in dividing cells, which is associated with a detectable increase in the levels and centrosomal accumulation of phosphorylated Rab10. Short-term application of a LRRK2 kinase inhibitor reverts the increase in phospho-Rab10, and this in turn leads to a rapid reversal of the centrosomal cohesion deficits. Such dynamic behaviour is consistent with the direct RILPL1-mediated recruitment of phospho-Rabs to a centrosomal location to cause an increase in the distance between duplicated centrosomes. It further predicts that the underlying mechanism may involve the dynamic recruitment and/or displacement of protein(s) necessary to keep duplicated centrosomes in close proximity to each other.

We have previously reported centrosomal cohesion deficits in lymphoblastoid cell lines from G2019S LRRK2-PD patients as compared to healthy controls, and such deficits were also observed in a subset of sporadic PD patients (11, 12). In future experiments, it will be interesting to determine whether centrosomal deficits can be detected in peripheral cells from PD patients harboring LRRK2 risk variants or mutations in vps35. In addition, our study shows that expression of N2081D LRRK2 mutant causes kinase activity-mediated centrosomal cohesion deficits. Since the N2081D LRRK2 mutation confers risk for PD as well as for Crohn’s disease (28), further studies are warranted to probe for centrosomal deficits in peripheral cells from Crohn’s disease patients, as this may aid in stratifying patients benefitting from LRRK2 kinase inhibitor therapeutics in clinical studies.

Recent work has shown that inducing lysosomal damage causes recruitment of LRRK2 to lysosomes, followed by the lysosomal accumulation of phospho-Rab10 (39–42). Conversely, mitochondrial depolarization causes the mitochondrial accumulation of Rab10 to facilitate mitophagy, and such accumulation is impaired in the context of pathogenic LRRK2 (43). Our studies were performed in the absence of treatments to induce lysosomal damage or mitochondrial depolarization. Under such normal physiological conditions, we find that the majority of phospho-Rab10 localizes to the centrosome to cause centrosomal cohesion and ciliogenesis deficits. In future experiments, it will be interesting to determine how triggers which lead to lysosomal or mitochondrial damage might impact upon the centrosomal deficits as described here. In either case, our data indicate that it is the RILPL1-mediated localization of phospho-Rab10 to the centrosome which is responsible for the cohesion deficits mediated by pathogenic LRRK2. Interestingly, the expression of a C-terminal fragment of RILPL1 which localizes to cytosolic punctate structures reverts the cohesion deficits mediated by pathogenic LRRK2 by redistributing phospho-Rab10 from its centrosomal location to those structures, without altering the total levels of phospho-Rab10 as assessed by Western blot analysis. Therefore, the subcellular location of the phospho-Rab10 accumulation, rather than total phospho-Rab10 levels, are relevant for our understanding of pathogenic LRRK2 action in a given cellular context.

Whilst the centrosomal cohesion and ciliogenesis deficits are a direct consequence of enhanced LRRK2 kinase activity and mediated by an increase in centrosomal phospho-Rab10 levels, the pathophysiological relevance of those alterations requires further investigation. Ciliogenesis deficits in cholinergic neurons in the striatum may lead to a loss of neuroprotective signaling onto dopaminergic neurons in the substantia nigra (9), but direct evidence for a lack of such neurotrophic support circuitry in the presence of pathogenic LRRK2 is currently missing. Similarly, ciliogenesis deficits in striatal astrocytes may dysregulate striatal neuronal circuitry (38), but further work is required to determine possible alterations in neuronal excitability in the striatum, and its effects on dopaminergic cell survival in the substantia nigra. Finally, the potential pathophysiological relevance for the LRRK2-mediated centrosomal cohesion deficits as observed in dividing cells remains unknown. Future experiments will seek to determine whether centrosomal cohesion deficits cause alterations in cell proliferation leading to impaired adult neurogenesis as described in transgenic G2019S LRRK2 mice (44), with potential relevance for various non-motor symptoms associated with PD.

Altogether, our data demonstrate that pathogenic LRRK2 mutations, LRRK2 risk variants and modulators of the LRRK2 signaling pathway all converge upon causing centrosomal cohesion and ciliogenesis defects. These deficits are dependent on RILPL1, and directly mediated by the centrosomal accumulation of phospho-Rab10. The localization of RILPL1 implicates the subdistal appendages of the mother centriole as the prime site of action for the LRRK2-mediated phospho-Rab10 accumulation, with downstream effects on centrosomal cohesion and ciliogenesis as described here.

## Materials and Methods

### DNA constructs and site-directed mutagenesis

GFP-tagged human wildtype LRRK2, pathogenic mutant LRRK2 (G2019S, R1441C, R1441G, Y1699C, N1437H, I2020T) kinase-inactive mutant LRRK2 (K1906M), as well as all mutant LRRK2 constructs containing R1398H, R1398L, T1343V, or R1398L/T1343V have been previously described (10, 11, 33). The T1410A, R1628P, S1647T, N2081D and G2385R mutant GFP-tagged LRRK2 constructs were generated by site-directed mutagenesis (QuikChange, Stratagene), and identity of all constructs verified by sequencing of the entire coding region. Flag-tagged human wildtype and mutant LRRK2 constructs, as well as N-terminally or C-terminally GFP-tagged human RILPL1 constructs have been previously described (10). RILPL1-miniSOG-HA construct was generated using Gibson Assembly Master Mix (New England Biolabs), and identity of construct verified by sequencing of the entire coding region. N-terminally HA-tagged human vps35, vps35-D620N, vps35-L774M, vps35-M57I, and C-terminal half of human RILPL1 [E280-end] tagged at the C-terminus with eGFP (RL1d-GFP) (4) were generous gifts from D. Alessi (University of Dundee, UK), and are available at https://mrcppureagents.dundee.ac.uk/. For transient transfections of mammalian cells, all DNA constructs were prepared from bacterial cultures grown at 37 °C using PureYield™ Plasmid Midiprep System (Promega) according to manufacturer’s instructions.

### Cell culture and transfections

HEK293T cells were cultured in full medium (Dulbecco’s modified Eagle’s medium (DMEM) containing low glucose and 10 % fetal bovine serum, non-essential amino acids, 100 U/ml penicillin and 100 μg/ml streptomycin) and transfected at 80 % confluence with 1 μg of LRRK2 constructs (and 100 ng of HA-tagged vps35 constructs where indicated) and 3 μl of LipoD293™ Transfection Reagent (SignaGen Laboratories) per well of a 12-well plate overnight. The next day, cells were split to 25 % confluence onto poly-L-lysine-coated coverslips and subjected to immunocytochemistry or Western blot analysis 48 h after transfection.

The various wildtype and CRISPR-Cas9 knockout A549 cells (Rab8a-KO, Rab10-KO, Rab12-KO, Rab35-KO, Rab43-KO, RILPL1-KO, RILPL2-KO, vps35-KO, Rab29-KO and PPM1H-KO) were generous gifts from D. Alessi (University of Dundee, UK) and have been previously described (4, 9, 14, 20, 21, 45, 46). Cells were cultured in DMEM containing high glucose without glutamine, and supplemented with 10 % fetal bovine serum, 2 mM L-glutamine, 100 U/ml of penicillin and 100 μg/ml of streptomycin. Cells were subcultured at a ratio of 1:6 - 1:10 twice a week and transfected at 90 % confluence. Cells were either transfected with 1 μg of LRRK2 constructs, co-transfected with 1 μg of LRRK2 and 100 ng RL1d-GFP construct, or co-transfected with 1 μg of pCMV and 100 ng of GFP-RILPL1, RILPL1-GFP or RILPL1-miniSOG constructs, along with 4 μl of LipoD293™ Transfection Reagent (SignaGen Laboratories) per well of a 12-well plate. Transfection media was replaced with full media after 5 h, cells were split 1:4 the next day and processed for immunocytochemistry or Western blotting 48 h after transfection.

In all cases, cells were grown at 37 °C and 5 % CO_2_ in a humidified atmosphere, and all lines were regularly tested for mycoplasma contamination. Where indicated, cells were treated with MLi2 (MRC PPU, Dundee, UK), or with the equivalent volume of DMSO before fixation.

### Immunocytochemistry

For centrosome staining, HEK293T and A549 cells were fixed with 4 % paraformaldehye (PFA) in PBS for 15 min at room temperature, followed by permeabilization with 0.2 % Triton-X100/PBS for 10 min at room temperature. Coverslips were incubated in blocking solution (0.5 % BSA (w/v) in 0.2 % Triton-X100/PBS) for 1 h at room temperature and incubated with primary antibodies in blocking solution overnight at 4 °C. Primary antibodies included rabbit polyclonal anti-pericentrin (Abcam, ab4448, 1:1000), mouse monoclonal anti-flag (Sigma, clone M2, F1804, 1:500), rat monoclonal anti-HA (Sigma, 11867423001, 1:500), mouse monoclonal anti-γ-tubulin (Abcam, ab11316, 1:1000), rabbit polyclonal anti-RILPL1 (Sigma, HPA-041314, 1:300), sheep polyclonal anti-RILPL1 (1:50), rabbit monoclonal knockout-validated anti-pT73-Rab10 (Abcam, ab241060, 1:1000), or rabbit polyclonal anti-pT72-Rab8a (1:500, generous gift of D. Alessi). Immunocytochemistry employing sheep antibodies was performed sequentially, with the sheep antibody employed first. Upon overnight incubation with antibodies, coverslips were washed twice with 0.2 % Triton-X100/PBS (wash buffer), followed by incubation with secondary antibodies in wash buffer for 1 h at room temperature. Secondary antibodies included Alexa405-conjugated goat anti-mouse, Alexa488-conjugated goat anti-mouse, goat anti-rabbit or donkey anti-sheep, Alexa555-conjugated goat-anti-mouse, goat anti-rabbit or donkey anti-sheep, and Alexa647-conjugated goat anti-mouse, goat anti-rabbit or donkey anti-sheep (all from Invitrogen, 1:1000). Coverslips were washed twice in wash buffer, rinsed in PBS and mounted in mounting medium with DAPI (Vector Laboratories).

### Image acquisition and quantification

Images were acquired on a Leica TCS-SP5 confocal microscope using a 63x 1.4 NA oil UV objective (HCX PLAPO CS) (10), or on an Olympus FV1000 Fluoview confocal microscope using a 60x 1.2 NA water objective lens. Images were collected using single excitation for each wavelength separately and dependent on secondary antibodies, and the same laser intensity settings and exposure times were used for image acquisition of individual experiments to be quantified. Around 10-15 optical sections of selected areas were acquired with a step size of 0.5 μm, and maximum intensity projections of z-stack images analyzed and processed using Leica Applied Systems (LAS AF6000) image acquisition software or ImageJ. Duplicated centrosomes were scored as being split when the distance between their centres was > 1.5 μm for HEK293T cells (10, 11) or > 2.5 μm for A549 cells (10), respectively. Mitotic cells were excluded from the analysis in all cases. Quantification of centrosomal distances was performed by an additional observer blind to condition, with identical results obtained in both cases.

### Electron microscopy, sample preparation and imaging

A549 cells were cultured in glass-bottom MatTek dishes (MatTek Life Sciences, P35G-0-14-C) and transfected with RILPL1-miniSOG-HA using LipoD293™ Transfection Reagent (SignaGen Laboratories, SL100668) as described above. Proteins were allowed to express for 48 hours and cells were processed as previously described (37, 47). Briefly, cells were rinsed with pre-warmed HBSS and fixed using pre-warmed 2.5 % (w/v) glutaraldehyde (Electron Microscopy Sciences, 16220), 0.1 % tannic acid (w/v) (Electron Microscopy Sciences, 21700), 3 mM calcium chloride in 0.1 M sodium cacodylate buffer pH 7.4 (Ted Pella Incorporated, 18851) for 5 minutes at 37 °C and then on ice for 1 hour. Subsequent steps were performed on ice, cells were rinsed 5 times using chilled 0.1 M sodium cacodylate buffer pH 7.4 (wash buffer) and treated for 30 minutes in a blocking solution (50 mM glycine, 10 mM KCN, 20 mM aminotriazole and 0.01 % hydrogen peroxide in 0.1 M sodium cacodylate buffer pH 7.4) to reduce non-specific background precipitation of DAB.

Cells were first imaged with minimum light exposure to identify transfected cells for Correlative Light and Electron Microscopy (CLEM) using a Leica SPE II inverted confocal microscope outfitted with a stage chilled to 4 °C. For photo -oxidation, DAB (3-3’-diaminobenzidine, Sigma-Aldrich, D8001-10G) was dissolved in 0.1 N HCl at a concentration of 5.4 mg/ml and subsequently diluted ten fold into blocking solution, mixed, and passed through a 0.22 μm syringe filter before use. DAB solution was freshly prepared prior to photo-oxidation and placed on ice protected from light. DAB solution was added to the MatTek dish and regions of interest were illuminated through a standard FITC filter set (EX470/40, DM510, BA520) with intense light from a 150 W Xenon lamp. Photo-oxidation was stopped as soon as an optically-dense brown reaction product began to appear in place of the miniSOG intrinsic green fluorescence signal, monitored by transmitted light (around 4-6 minutes). Multiple areas on a single MatTek dish were photo-oxidized.

Subsequently, plates were placed on ice and washed 5 times 2 minutes each with ice-cold wash buffer to remove unpolymerized DAB. After washing out DAB, cells were post-fixed with 2 % reduced osmium tetroxide (Electron Microscopy Sciences, 19190) (2 % osmium tetroxide, 1.5 % KFeCN in 0.1 M sodium cacodylate buffer pH 7.4) for 1 hour on ice, then washed with ice-cold double-distilled water 3 times for 1 minute. Some samples were additionally stained overnight with filtered 2 % uranyl acetate (Electron Microscopy Sciences, 22400) in double-distilled water and compared to others in which this step was omitted. The following day, plates were washed 3 times for 1 minute with double-distilled water and were dehydrated with an ice-cold graded ethanol series (20 %, 50 %, 70 %, 90 %, 100 %, 100 %, 3 minutes each) and washed once at room temperature anhydrous ethanol (3 minutes). Samples were then embedded in Durcupan™ ACM resin (Sigma Aldrich; Durcupan™ ACM component A, M epoxy resin (44611); Durcupan™ ACM component B, hardener 964 (44612); Durcupan™ ACM component C, accelerator 960 (44613); Durcupan™ ACM component D (44614)) using a 1:1 mixture of anhydrous ethanol:Durcupan™ ACM resin for 30 minutes on a platform with gentle rocking, followed by incubation with 100 % Durcupan™ ACM resin overnight with rocking. The following day, the resin was removed from MatTek dishes by decanting and gentle scraping without touching the cells and changed with freshly prepared resin for 1 hour (three times). After third replacement, resin was polymerized in a vacuum oven at 60 °C for 48 hours under - 10 mm Hg vacuum pressure atmosphere.

Photo-oxidized areas of interest were identified by transmitted light, sawed out using a jeweller’s saw, and mounted on dummy acrylic blocks with cyanoacrylic adhesive. The coverslip was carefully removed, the resin was trimmed, and ultrathin sections (80 nm thick) were cut using a diamond knife (Diatome). Electron micrographs were recorded using a FEI Tecnai™ 12 Spirit TEM (transmission electron microscope) operated at 80 kV. For electron tomography, thicker sections (750 nm) were imaged on a FEI Titan Halo™ microscope operated at 300 kV in scanning mode, the scattered electrons being collected on a high angle annular dark field detector. Prior to imaging, the luxel grids carrying the specimen serial sections were coated with carbon on both sides; colloidal gold particles (10, 20 and 50 nm diameter) were deposited on each side of the sections to serve as fiducial markers. Because centrosomes cannot be clearly distinguished on electron micrographs of thick sections, a preliminary tomography run was first implemented using a low magnification setting on the cells of interest (spanning a 12 μm x 12 μm area). This allowed to pinpoint the exact sections containing centrosomal areas; higher resolution tomograms (with a ~1 nm pixel size) were then acquired on the spot. For each tomogram, four tilt series were collected using the SerialEM package. For each series, the sample was tilted from −60 to +60 degrees, every 0.5 degree. Tomograms were generated using an iterative reconstruction procedure (48).

### Western blotting

HEK293T and A549 cells were collected 48 h after transfection from a well of a 6-well plate. Cells were washed in PBS, and the cell pellet was resuspended in 75 μl PBS. Cells were lysed with 25 μl of 4x Nu-PAGE LDS sample buffer (Novex, Life Technologies, NP00008) supplemented with β-mercaptoethanol to a final volume of 2.5 % (v/v), sonicated and boiled at 70 °C for 10 min. Around 10-15 μl (around 20 μg of protein) were resolved by SDS-PAGE polyacrylamide gel electrophoresis using 4-20 % precast gradient gels (Bio-Rad, 456-1096), and proteins were electrophoretically transfected onto nitrocellulose membranes (GE Healthcare). Membranes were blocked in blocking buffer (Li-COR Biosciences, Li-COR Odyssey intercept blocking buffer, 927-70001) for 1 h at room temperature, and incubated with primary antibodies in blocking buffer overnight at 4 °C. Primary antibodies included mouse monoclonal anti-GFP (Sigma, 11814460001, 1:1000), mouse monoclonal anti-flag (Sigma, clone M2, F1894, 1:500), rat monoclonal anti-HA (Sigma, 118674123001, 1:500), mouse monoclonal anti-Rab8 (BD, 610844, 1:500), mouse monoclonal knockout-validated anti-Rab10 (Sigma, SAB5300028, 1:1000), rabbit monoclonal anti-pT72-Rab8a (Abcam, ab230260, 1:1000), rabbit monoclonal anti-pT73-Rab10 (Abcam, ab230261, 1:1000), rabbit monoclonal anti-S935-LRRK2 (Abcam, ab133450, 1:500), mouse monoclonal anti-α-tubulin (Sigma, clone DM1A, T6199, 1:25’000) and mouse monoclonal anti-GAPDH (Santa Cruz, sc-32233, 1:2000). Sheep polyclonal antibodies against Rab12, Rab35 and Rab43 (generous gifts from D. Alessi) were employed at 1:100 dilution in 5 % milk in 0.1 % Tween-20/TBS blocking buffer overnight at 4 °C, and membranes were developed using ECL Prime Western Blotting Detection Reagent (GE Healthcare) as previously described (10).

### Statistical analysis

One-way ANOVA with Tukey’s *post hoc* test was employed, with significance set at p < 0.05. Significance values for all data are indicated in the figure legends. All statistical analysis and graphs were performed with Prism software version 7.0 (GraphPad, San Diego, CA).

## Acknowledgements

We are especially grateful to Dario Alessi for providing a large variety of reagents. We thank Laura Montosa and Hiroyuki Hakozaki (UCSD) for technical assistance with confocal microscopy, and Mason Mackey (UCSD) for technical assistance with electron microscopy. We also thank Junru Hu for lab assistance and Andrea Thor for help with EM sample preparation.

