## Supplemental Figure Legends for "LRRK2 causes centrosomal deficits via phosphorylated Rab10 and RILPL1 at centriolar subdistal appendages"

**Figure S1.** The protective R1398H variant decreases centrosomal cohesion deficits mediated by pathogenic LRRK2. (**A**) HEK293T cells were transfected with GFP-tagged wildtype (wt) LRRK2, pathogenic G2019S mutant, or pathogenic G2019S mutant containing the protective R1398H variant (G2019S-RH). Cells were stained for the centrosomal marker pericentrin (red) and DAPI (blue). Arrows point to centrosomes in transfected cells. Scale bar, 10 μm. (**B**) Quantification of the percentage of cells with split centrosomes (duplicated centrosomes with a distance between their centres > 1.5 μm) from either non-transfected cells, or from cells transfected with the indicated constructs in the absence or presence of MLi2 (100 nM, 2 h) as indicated. Bars represent mean ± S.E.M. (n=3 experiments); ****p < 0.001; ***p < 0.005.

**Figure S2.** Synthetic variants which modulate GTP binding/hydrolysis decrease centrosomal cohesion deficits mediated by pathogenic LRRK2. (**A**) HEK293T cells were transfected with GFP-tagged wildtype (wt) LRRK2, pathogenic G2019S mutant, pathogenic G2019S mutant containing the synthetic R1398L variant (G2019S-RL), the T1343V mutation (G2019S-TV), or both (G2019S-RLTV). Cells were stained for the centrosomal marker pericentrin (red) and DAPI (blue). Arrows point to centrosomes in transfected cells. Scale bar, 10 μm. (**B**) Quantification of the percentage of cells with duplicated split centrosomes from either non-transfected cells, or from cells transfected with the indicated constructs in the absence or presence of MLi2 (100 nM, 2 h) as indicated. Bars represent mean ± S.E.M. (n=3 experiments); ****p < 0.001; ***p < 0.005.

**Figure S3.** Pathogenic LRRK2-mediated centrosomal cohesion deficits are blunted in Rab29-KO, but not in vps35-KO cells. (**A**) Example of A549 Rab29-KO cells transfected with either flag-tagged wildtype (wt) or Y1699C-mutant LRRK2, and either treated or untreated with 500 nM MLi2 for 2 h prior to immunocytochemistry with antibodies against flag, pericentrin and with DAPI. Scale bar, 10 μm. (**B**) Quantification of the percentage of A549 wildtype cells, A549-Rab29KO cells or A549-vps35-KO cells with split centrosomes either in the absence or transfection (ctrl), or upon transfection with either wildtype or Y1699C-mutant LRRK2. Cells were treated with or without MLi2 (500 nM, 2 h) prior to immunocytochemistry as indicated. Bars represent mean ± S.E.M. (n=3-5 independent experiments); ****p < 0.001; ***p < 0.005.

**Figure S4.** Localization of phospho-Rab10 is influenced by the presence of RILPL1. Examples of wildtype, RILPL1-KO or RILPL2-KO A549 cells transfected with flag-tagged Y1699C LRRK2, and treated with or without MLi2 (500 nM, 2 h) before immunostaining with an antibody against phospho-Rab10 (p-Rab10) and with DAPI. Perinuclear phospho-Rab10 clusters are prominent in A549 wildtype cells expressing pathogenic LRRK2. In RILPL1-KO cells, perinuclear clusters are rarely observed, but phospho-Rab10 displays an additional punctate staining throughout the cytosol. In RILPL2-KO cells, perinuclear phospho-Rab10 staining is prominent in some cells expressing pathogenic LRRK2. Scale bar, 10 μm.

**Figure S5.** Endogenous RILPL1 localizes to centrosome and recruits phospho-Rabs in HEK293T cells transfected with pathogenic LRRK2. (**A**) Example of HEK293T cells transfected with GFP-tagged wildtype or pathogenic Y1699C LRRK2 and stained with antibodies against pericentrin (Alexa-647 secondary antibody, red), RILPL1 (Alexa-594 secondary antibody, pseudocolored in blue) and with DAPI (cyan). Arrows point to colocalization of endogenous RILPL1 with the centrosomal marker pericentrin. Scale bar, 10 μm. (**B**) Example of non-transfected cells (ctrl), or cells transfected with GFP-tagged wildtype or Y1699C LRRK2 and stained with antibodies against RILPL1 (Alexa-594 secondary antibody, red), phospho-Rab8 (Alexa-405 secondary antibody, pseudocolored in blue) and with TOPRO (cyan). The phospho-Rab8 antibody detects both phospho-Rab8 and phospho-Rab10 by immunocytochemistry (Lara Ordóñez et al., 2019). Arrow points to pathogenic LRRK2-expressing cell, where phospho-Rab accumulation colocalizes with endogenous RILPL1. Scale bar, 10 μm.
