## Supplementary figures and images for "LRRK2 causes centrosomal deficits via phosphorylated Rab10 and RILPL1 at centriolar subdistal appendages"

### Supplemental Figures

#### Slide 1
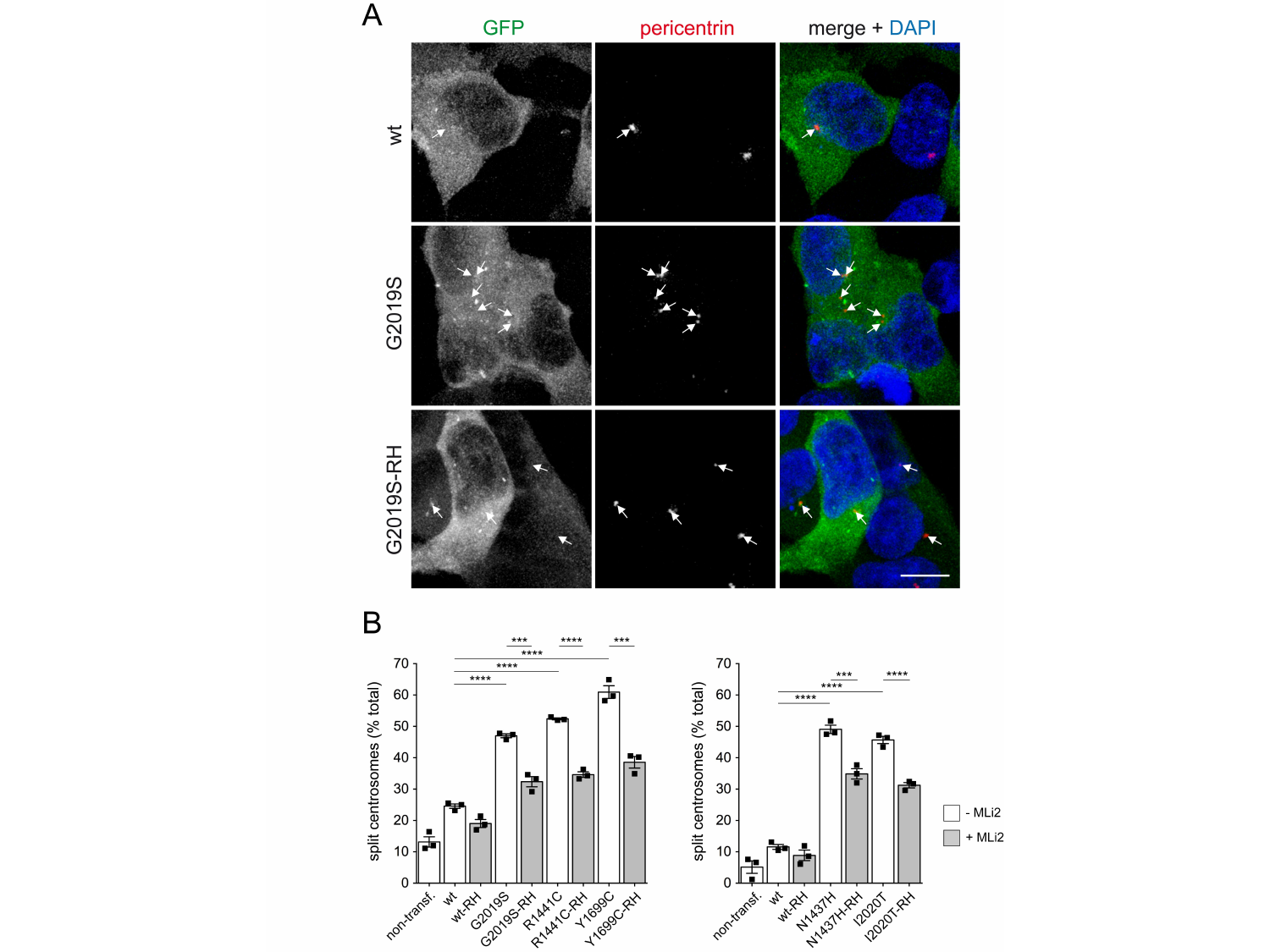

#### Slide 2
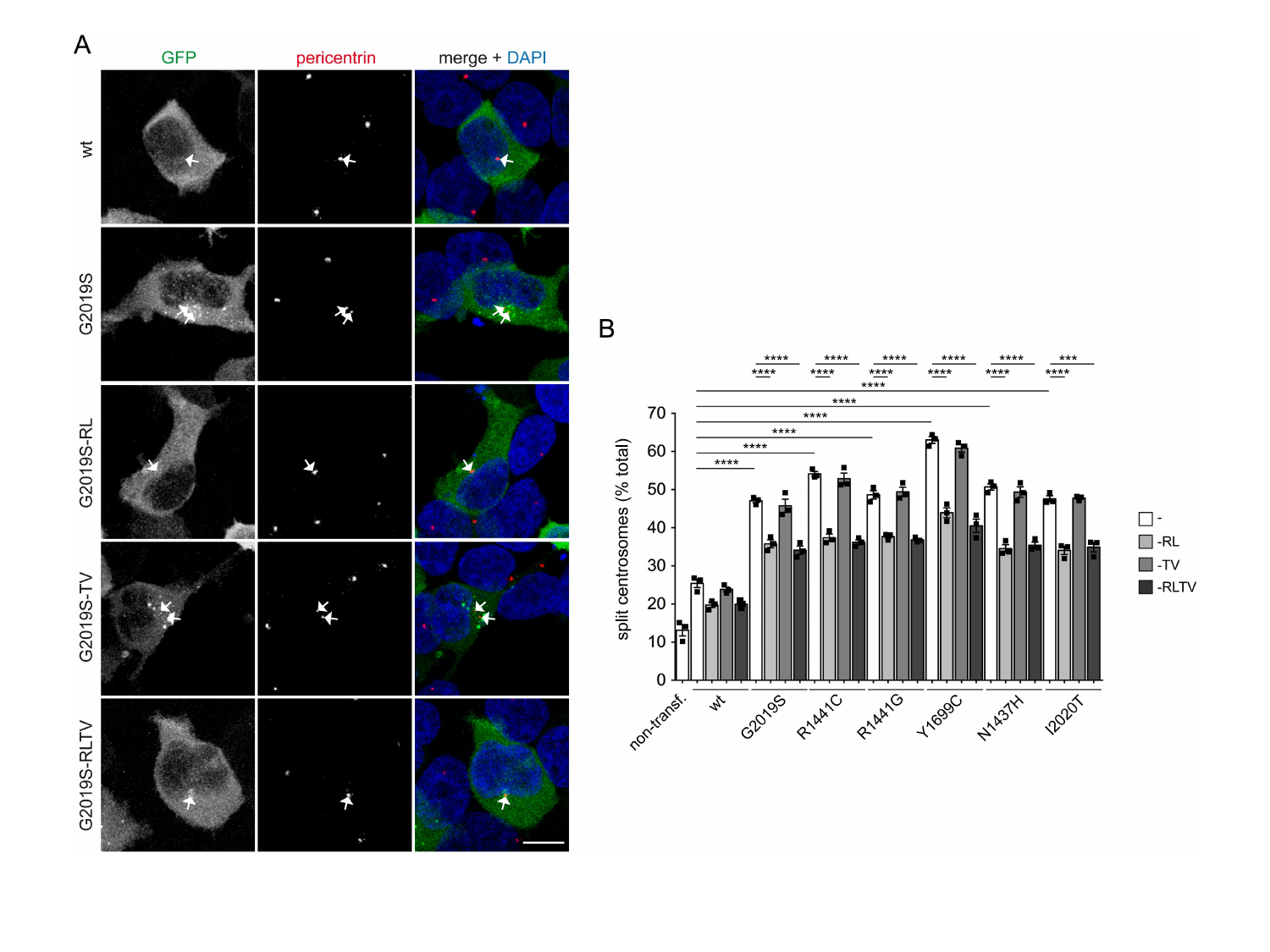

#### Slide 3
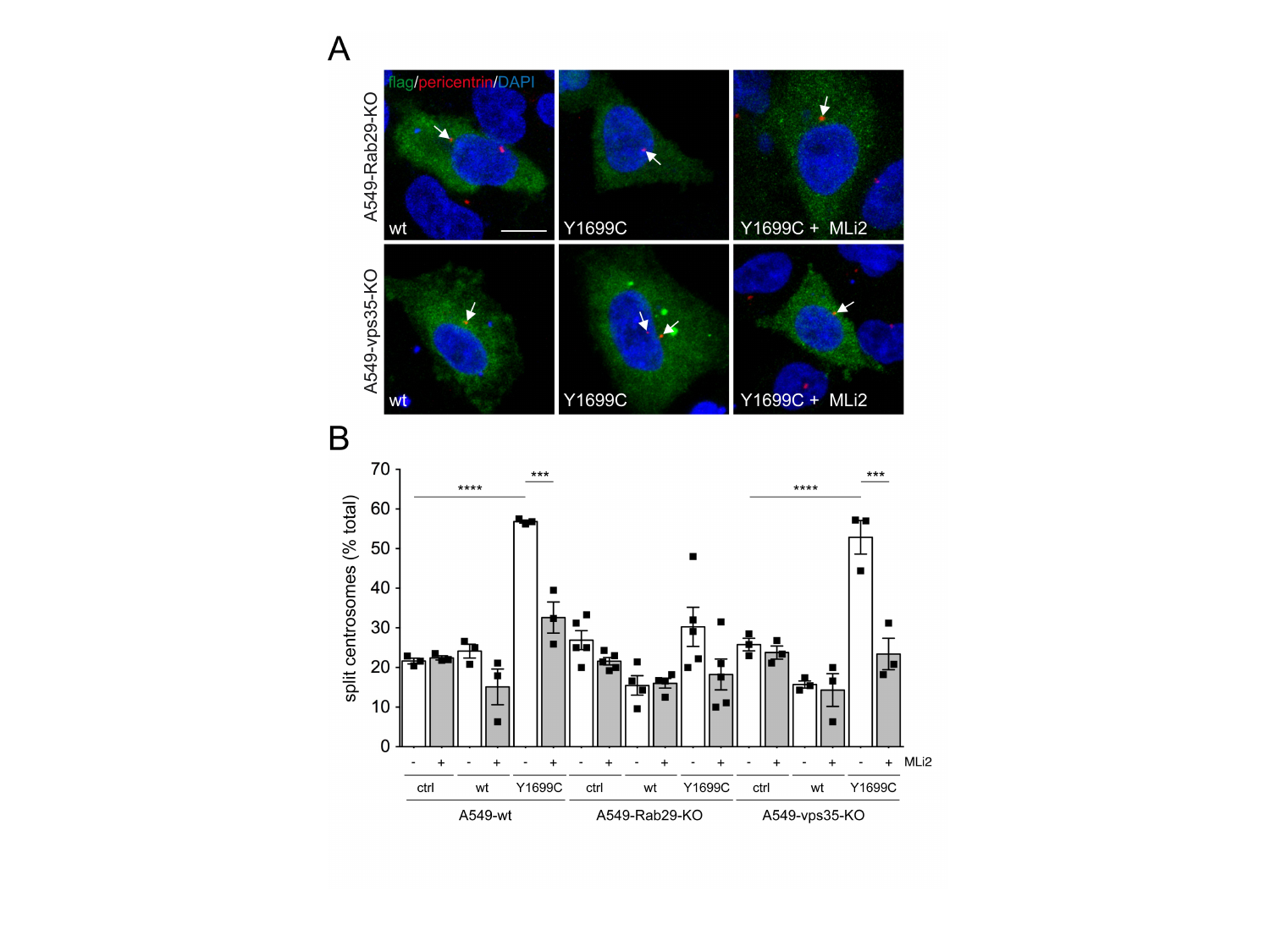

#### Slide 4
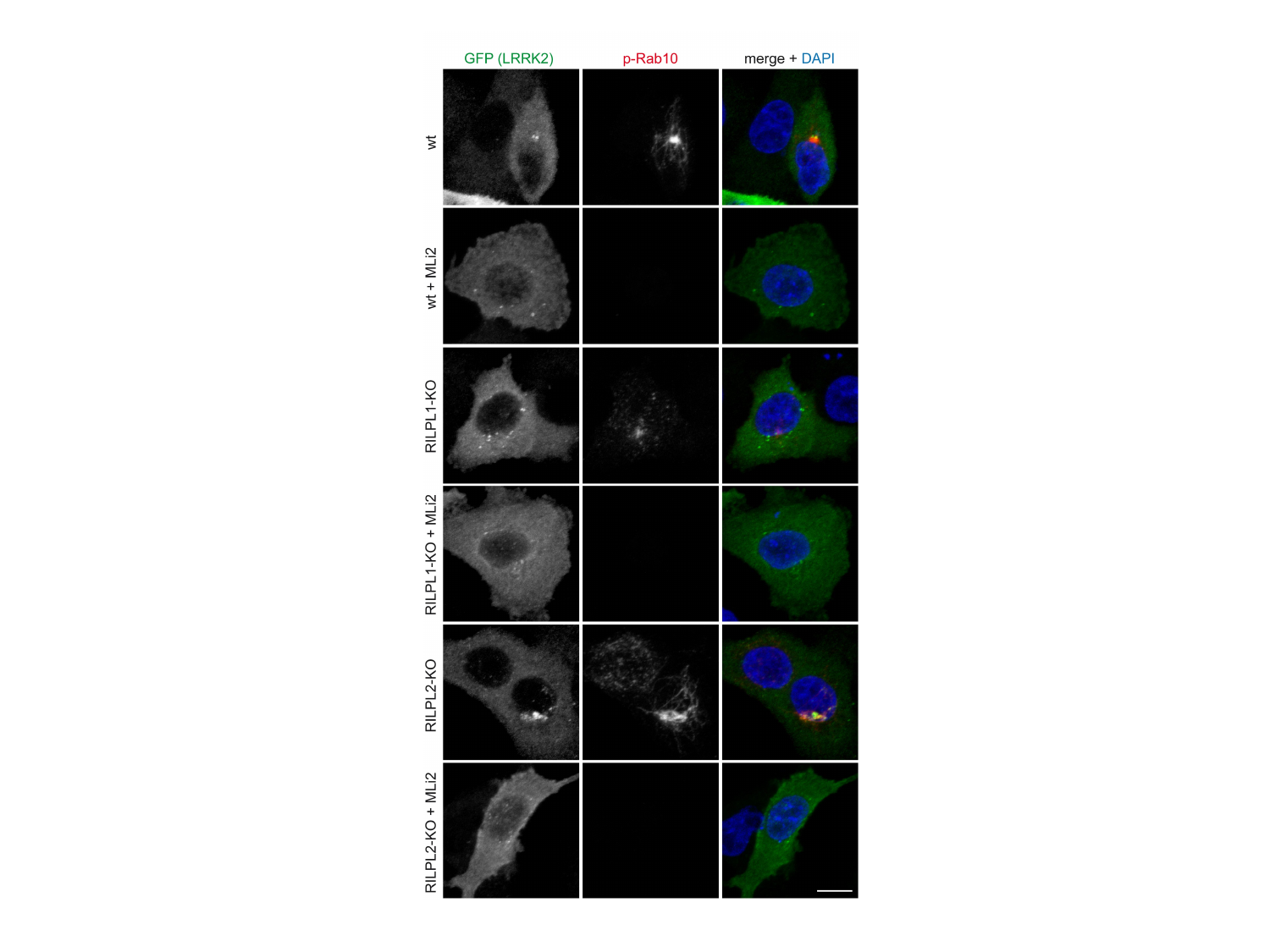

#### Slide 5
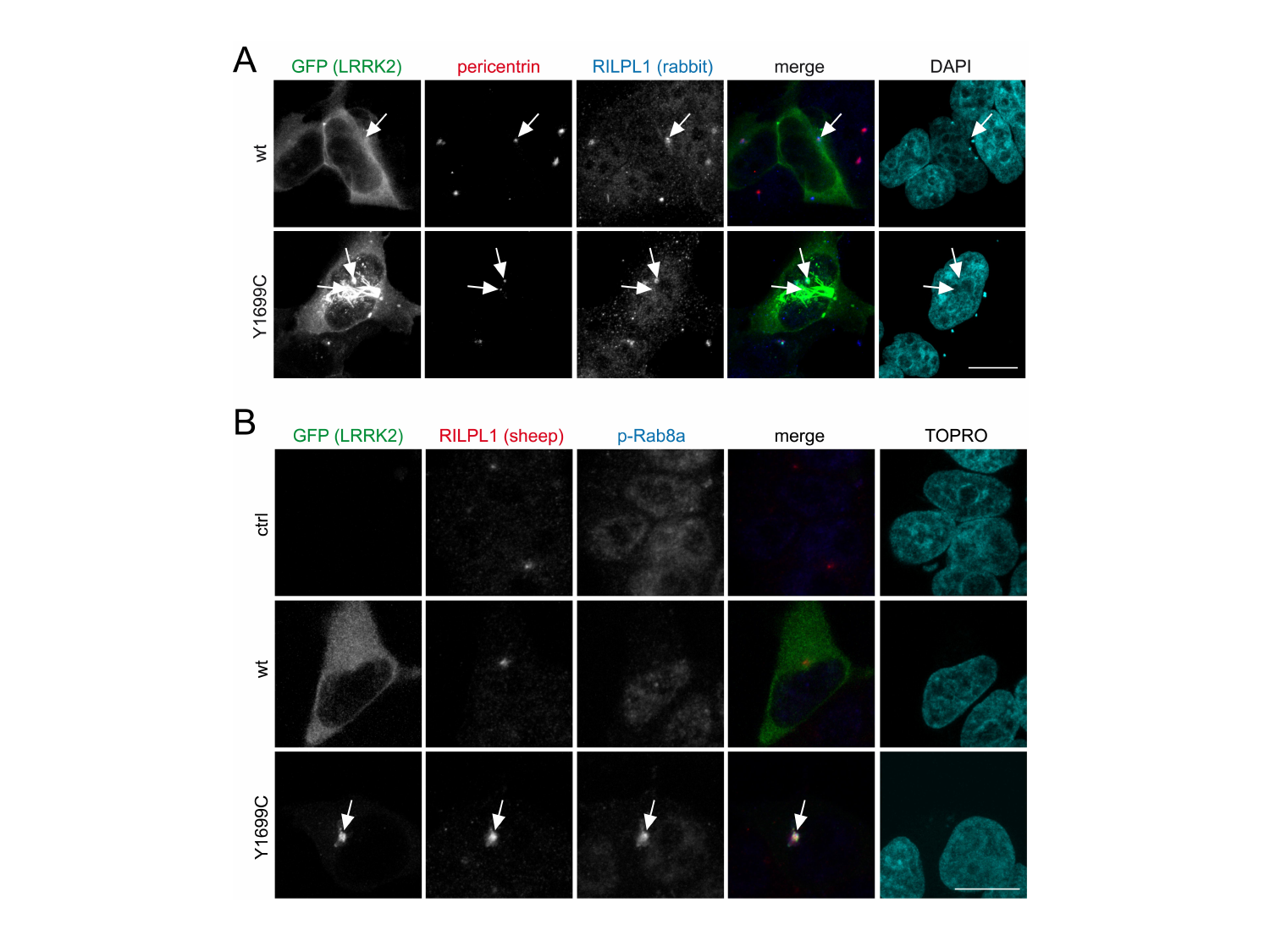
